# Microbial lysates as low-cost serum replacements in cellular agriculture media formulation

**DOI:** 10.1101/2024.10.28.620756

**Authors:** James Dolgin, Damayanti Chakravarty, Sean F. Sullivan, Yiming Cai, Taehwan Lim, Pomaikaimaikalani Yamaguchi, Joseph E. Balkan, Licheng Xu, Aaron D. Olawoyin, Kyongbum Lee, David L. Kaplan, Nikhil U. Nair

## Abstract

Cultivated meat, the process of generating meat in vitro without sacrificing animals, is a promising alternative to the traditional practice of livestock agriculture. However, the success of this field depends on finding sustainable and economical replacements for animal-derived and expensive fetal bovine serum (FBS) that is typically used in cell culture processes. Here, we outline an effective screening process to vet the suitability of microbial lysates to support the growth of immortalized bovine satellite cells (iBSCs) and mackerel (Mack1) cells. We show that easily producible, low-cost whole-cell lysates from *Vibrio natriegens* can be used to create serum-free media for the long-term growth of iBSCs. The optimized medium, named “VN40” (basal B8 media containing Vibrio natriegens lysate proteins at 40 µg/mL), outperforms previously established serum-free media while maintaining cell phenotype and myogenicity. Overall, this study shows a novel approach to producing serum-free media for cultivated meat production using microbially-derived lysates.

## 1. INTRODUCTION

The environmental impact of meat consumption is expected to continue to rise with increasing demand and the increasing global population (OECD, Food, & Nations, 2021). The production of beef using conventional livestock agriculture is unsustainable due to greenhouse gas emissions, eutrophication, deforestation, overuse of water, and the spread of zoonotic disease (Hayek, 2022; Poore & Nemecek, 2018). Similarly, modern fishing practices have resulted in 90% of the world’s fish stocks to be overexploited or collapse (Boyd et al., 2020; Grimaldo et al., 2023).

Cultivated meat – meat products generated from *in vitro* cultures of animal cells – represents a potentially sustainable alternative to conventional meat and fish production, which can reduce negative externalities and benefit animal welfare (Sinke et al. 2023; Treich, 2021). While conventional meat is responsible for 15% of global greenhouse gas emissions, cultivated meat can reduce this almost 13-fold (Sinke et al., 2023), especially through the use of renewable energy sources. Much progress has been made recently in developing stable immortalized lines of both bovine (Stout et al., 2023a) and mackerel (Saad et al., 2023) muscle satellite cells, leading to the creation of “iBSC” (Immortalized Bovine Satellite Cell) and “Mack1” cell lines, respectively. However, mammalian and marine cell cultures still largely depend on the use of fetal bovine serum (FBS), an expensive and animal-based ingredient with significant lot-to-lot variability (Zheng et al., 2006) and risks of contamination factors (Urz et al., 2022), making its use in cultivated meat infeasible. With 90% of the cost of cultivated meat production stemming from growth media (Specht, 2020), inexpensive FBS alternatives are needed to bring cultivated meat towards commercial production. Many attempts have been made to create serum-free media (SFM) for cultivated beef and fish with varying levels of success. Of note is recent work adapting serum-free “B8” media designed for human iPSC culture (Kuo et al., 2020) to the culture of bovine satellite cells (Stout et al., 2022) via the addition of recombinant albumin (rAlbumin), leading to the creation of “Beefy 9” medium. Still, efforts to replace FBS with recombinant albumin (Kolkmann, Van Essen, Post, & Moutsatsou, 2022; Stout et al., 2022) or other mitogenic proteins like fetuin (Skrivergaard et al., 2023) for cultivated meat production require expensive purified recombinant protein on the g/L scale. Current estimates show cultivated meat may require as much as 42 liters of media per kilogram to produce (Robert Vergeer, 2021), making the dependence on such quantities of recombinant proteins economically infeasible. Other methodologies using crude extracts or hydrolysates from plant (Stout et al., 2023) or microalgal (Dong et al., 2024; Okamoto et al., 2022) sources to supplement SFM have the potential to reduce raw material costs considerably. However, these methods can be labor-intensive, (Stout et al., 2023b; Ho et al., 2021), require corrosive chemicals (Okamoto et al., 2022), or are otherwise still dependent on animal serum (Dong et al., 2024) or animal by-products (Andreassen et al., 2020).

Microbes such as yeast, bacteria, and algae may provide a promising solution to the longstanding problem of creating serum-free media formulations, as they are animal-free, inexpensive to produce, highly renewable, and rich in proteins and other nutrients. Microbes can also be engineered to express growth factors, which currently represent 99% of the cost of cultured meat media at scale (Specht, 2020). Yeast hydrolysates have longstanding use in creating serum-free media (Chevalot et al., 2010; Sung et al., 2004), and recent reports detail using extracts from *Saccharomyces cerevisiae* (Lei et al., 2023), *Enterococcus hirae* (Celebi-Birand et al., 2023), cyanobacteria (Ghosh et al., 2023; Jeong et al., 2021; Tuomisto & de Mattos, 2011), and *Chlorella* algae (Dong et al., 2023; Okamoto et al., 2022) in cultured meat production. However, current research has not demonstrated success in identifying a non-pathogenic, rapidly replicating microbial extract capable of stimulating long-term cellular growth in serum-free conditions. Microalgae are slow-growing, taking several days to culture (Okamoto et al., 2022), while cyanobacteria can have toxic effects on cells (Chia et al., 2018). Yeast and bacterial extracts have not been shown to completely substitute for FBS or support long-term growth of animal cells (Celebi-Birand et al., 2023; Chia et al., 2018). A more comprehensive list of previous attempts to replace serum in cultivated meat media is in **Table S1.**

Media to produce cultivated meat must strive to minimize time, labor, and material costs in order to make cultivated meat economically viable. Such media must also be animal-free and safe to consume and produce. Finally, cultivated meat media must stimulate long-term cell growth with minimal impacts on cell characteristics. While microbial extracts are a promising ingredient for replacing FBS in cell culture media, there have not been any results that satisfy the criteria necessary for commercial use in cultivated meat production. Further, it is known that media requirements vary significantly depending on cell type (O’Neill et al., 2022).

In this work, we tested multiple lysates from Gram-positive and Gram-negative bacteria, as well as a yeast, to determine their suitability as replacements for FBS in bovine and fish cell culture media. Based on an initial screen, we found that lysates derived from *Vibrio natriegens* could serve as an alternative to FBS for iBSCs. A similar screen also identified that lysates from *Saccharomyces cerevisiae* could support growth of Mack1 cells in reduced serum conditions. We show that iBSCs can be adapted to grow rapidly in *Vibrio natriegens* lysate-containing media by long-term and short-term serial passaging. Cells cultured in this microbial lysate media (called VN40) demonstrate robust growth, preserve their satellite cell phenotype, and the ability to differentiate into multinucleated myotubes. These results highlight the promise of the whole-cell lysate derived from the fast-growing non-pathogenic marine bacterium, *V. natriegens*, as an easily producible, low-cost supplement for long-term serum-free growth of iBSCs for cultivated meat. Further, our approach may be generalizable to help identify novel microbial lysates as FBS substituents for different cultivated meat cell lines.

## 2. MATERIAL & METHODS

### 2.1 Culture and extraction of microbial lysate

*Escherichia coli* BL21(DE3) and Nissle 1917 (“BL21” and “ECN” respectively), *Vibrio natriegens* ATCC14048 (“VN”), *Lactobacillus plantarum* WCFS1 (“LP”), *Bacillus subtilis* 168 (“BS”)*, Lactococcus lactis* MG1363 (“LL”) and *Saccharomyces cerevisiae* CEN.PK (“CEN.PK”) were streaked out on plates using their respective solid media (**Table 1**). Single colonies from the plates were used to inoculate 25 mL of overnight cultures. The next day, cells at stationary phase were harvested by centrifuging at 4000 rpm for 5 min (ThermoScientific Sorvall Legend X1R Centrifuge). Harvested cells were prepared for sonication by washing and resuspending in 1 mL sterile phosphate buffer saline (PBS), without any additives. To generate microbial lysates, cells were sonicated (Branson 150 sonicator and 10 s Pulse, 30 s gap between each pulse and a total 7 min at 45 % amplitude) on ice. After sonication, the soluble lysate was obtained by centrifuging the sonicated solution at 21000 × g for 20 min. The supernatant was then filtered through a 0.2 μm syringe filter (Corning, #431219). The total protein content of the lysate was determined using Bradford Assay (Coomasie Plus^TM^ Protein Assay Reagent, #1856210) (**Figure 1A**).

**Figure 1.**
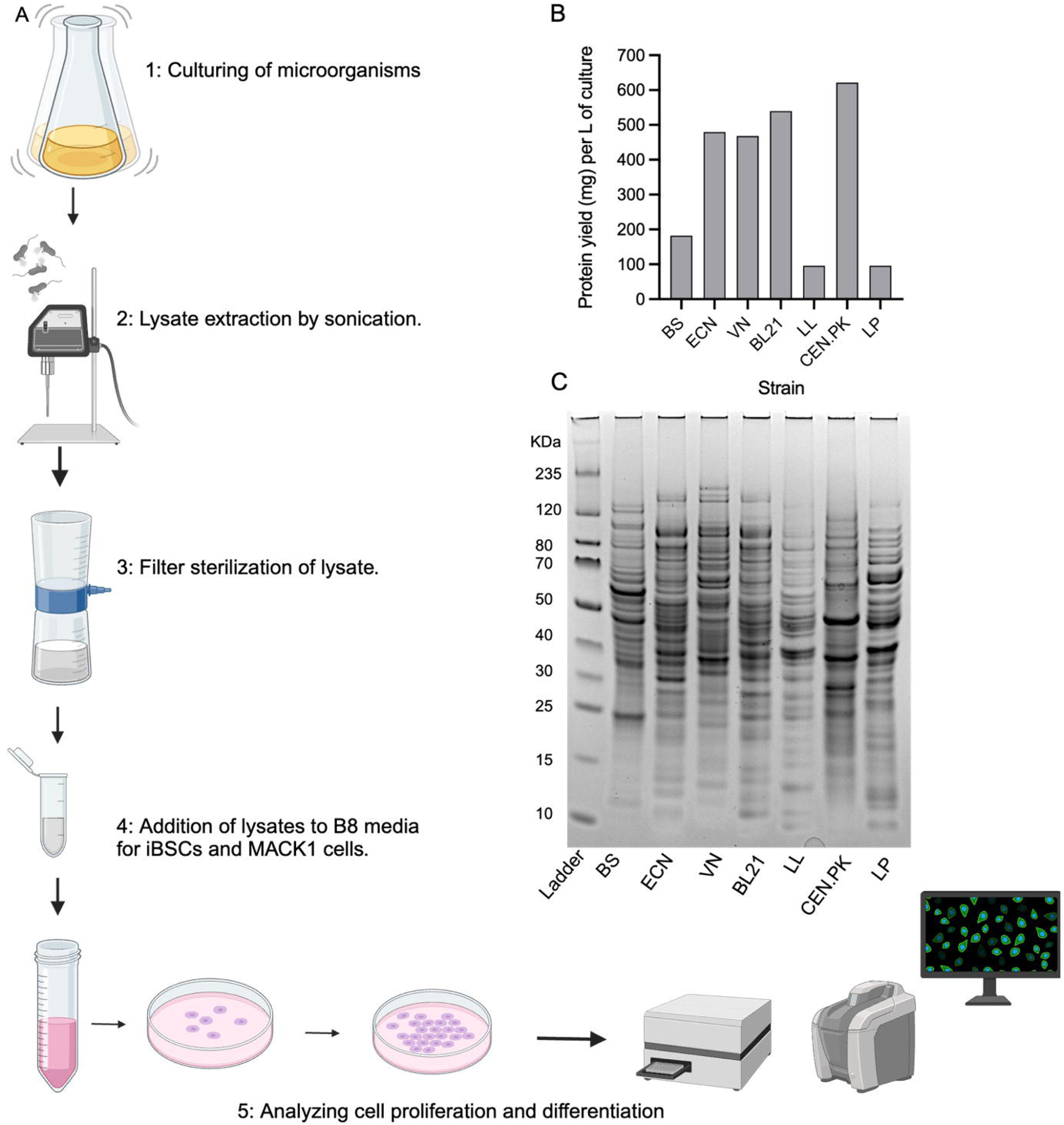
Workflow – screening of lysates from various microorganisms for muscle satellite cell performance. **A)** Overview of microbial workflow: microbial biomass production, lysate generation by mechanical lysis, and screening for proliferative activity on animal cells. After filter sterilization, the lysate is added to basal media, which is followed by estimating cell proliferation and differentiation. **B)** Higher protein yield resulted from Gram negative bacteria and yeast compared to gram positive bacteria. **C)** SDS-PAGE profile of seven lysates used in the initial screening. (L-R) Ladder, *B. subtilis* 168 (BS), *E. coli* Nissle 1917 (ECN), *V. natriegens* ATCC14048 (VN), *E. coli* BL21(DE3) (BL21), *L. lactis* MG1363 (LL), *S. cerevisiae* CEN.PK (CEN.PK), and *L. plantarum* WCFS1 (LP).

**Table 1.** Media and growth conditions for microbes cultivated for lysate generation.

| Microorganisms | Medium | Growth Conditions |
| --- | --- | --- |
| *E. coli* BL21(DE3) (BL21), *E. coli* Nissle 1917 (ECN), *Bacillus subtilis* 168 (BS) | Lysogeny Broth (LB) | Overnight at 37 °C |
| *Vibrio natriegens* ATCC14048 (VN) | LB with 5 % NaCl | Overnight at 37 °C |
| *Lactiplantibacillus plantarum* WCFS1 (LP), *Lactococcus Lactis* MG1363 (LL) | De Man–Rogosa–Sharpe (MRS) | Overnight at 37 °C |
| *S. cerevisiae* CEN.PK (CEN.PK) | Yeast peptone adenine dextrose (YPAD) | Overnight at 30 °C |

### 2.2 SDS-PAGE

Denaturing SDS-PAGE was performed following the Invitrogen NuPAGE® protocol. Lysate samples containing 30 µg of protein were adjusted to a final volume of 50 µL with PBS and mixed with 10 µL of 6× SDS loading buffer. The samples were incubated at 95 °C for 10 min. Subsequently, 10 µL of the sample was loaded onto a pre-cast NuPAGE Novex 4-12% Bis-Tris 1.0 mm mini-gel (Invitrogen). For each gel run, 5 µL of pre-stained SDS-PAGE standards (Bio-Rad) were included. Electrophoresis was conducted at a constant voltage (200 V) in 1× MES (2-morpholinoethanesulphonic acid) buffer solution at room temperature for approximately 45 min.

### 2.3 Lysate fractionation

*Vibrio natriegens* lysate was fractionated using a 3 kDa molecular weight cutoff filter (MilliporeSigma™ Amicon™ Ultra-15 Centrifugal Filter) at 4000 × g. This resulted in two fractions, i.e., protein rich fraction greater than 3 kDa (HMW fraction) and low molecular weight fraction lower than 3 kDa (LMW fraction). The fractions were then tested individually on iBSC cells. To test for additive effect of fractions, we used **Equation 1** and **Equation 2** to determine the sum of means (*µ*) and standard deviations (σ), respectively, of fractions’ impact on cell growth. These effects were compared to the impact of whole cell lysates, which are HMW and LMW fractions combined

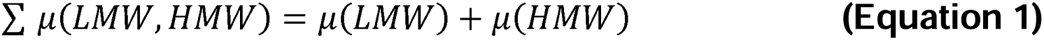

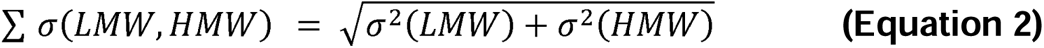

### 2.4 iBSC and Mack1 culture and maintenance

iBSCs were previously isolated from the semitendinosus of a 5-week-old Simmental calf at the Tufts Cummings School of Veterinary Medicine (Stout et al., 2022).These cells were engineered to constitutively express bovine telomerase reverse transcriptase (TERT) and cyclin-dependent kinase 4 (CDK4) in prior studies (Stout et al., 2023a). iBSCs at passage 40 were thawed from liquid nitrogen stock and passaged in iBSC growth media (iBSCGM) containing DMEM+Glutamax (ThermoFisher #10566024, Waltham, MA, USA) with 20 % FBS (ThermoFisher #26140079), 1 ng/mL human fibroblast growth factor 2 (FGF-2) (PeproTech #100-18B, Rocky Hill, NJ, USA), supplemented with 1 % antibiotic/antimycotic (ThermoFisher #1540062) and 2.5 μg/mL puromycin (ThermoFisher #A1113803) as a selection compound. iBSCs were cultured at 37 °C and 5 % humidity on tissue culture polystyrene (TCPS) and routinely passaged at 80–90 % confluency using 0.25 % trypsin-EDTA (ThermoFisher #25200056) to disassociate cells from TCPS. During passaging, cells were dissociated, centrifuged at 300 × g, and seeded at 1,500–2,000 cells/cm^2^. Mack1 cells (Kerafast, #ETU008-FP) isolated from Atlantic mackerel were cultured in Mack growth media (MackGM) containing Leiovitz’s L-15 medium (ThermoFisher #11415064) with 20 % FBS, 20 mM HEPES (Sigma Aldrich #H4034, St. Louis, MO, USA) buffer solution (pH 7.4), 1 % antibiotic/antimycotic, 1 ng/mL human FGF-2, and 10 µg/mL gentamicin (Sigma Aldrich #G1397)(Boyd et al., 2020; Saad et al., 2023). Mack1 cells were incubated at 27 °C without CO_2_ on TCPS, routinely detached at 70–80 % confluency using 0.05% trypsin-EDTA (ThermoFisher #25300054), centrifuged at 300 × g, and seeded at 3,500–4,000 cells/cm^2^. Both cell lines (iBSCs and Mack1) were fed every 2-3 days with iBSCGM and MackGM, respectively.

### 2.5 Short-term iBSC and Mack1 cell growth screening with microbial lysates

For iBSC short-term screening on microbial lysate-based media, iBSCs were harvested and resuspended into serum-free B8 media containing Hi-Def B8 aliquots (Defined Bioscience #LSS-204, San Diego, CA, USA) in DMEM/F12 (ThermoFisher #11320032) supplemented with 1 % antibiotic/antimycotic and 2.5 μg/mL puromycin. Cells were seeded onto 96-well plates at 2,500 cells/cm^2^ with 1.5 μg/cm^2^ of truncated recombinant human vitronectin (rhVTN-N; ThermoFisher #A31804). Cells were allowed to adhere for >3 h, then were washed once with DPBS (ThermoFisher #14190250) and media was changed to B8 + microbial lysates at given protein concentrations. Negative control conditions were fed B8 alone. Positive control conditions were fed Beefy 9, containing B8 supplemented with 0.8 mg/mL of recombinant albumin (rAlbumin, Sigma Aldrich #A9731)(Stout et al., 2022). Mack1 cells were harvested and resuspended in Mack1GM and plated onto 96-well plates at 2,500 cells/cm^2^. After more than 3 h incubation, cells were washed once with DPBS and media was changed to 2.5 % FBS-containing L-15 media (other media components, including HEPES, antibiotic/antimycotic, FGF-2, and gentamicin, were added at the same concentrations as Mack1GM) + microbial lysates at give protein concentrations. Negative controls were Mack1GM without FBS and 2.5% FBS-containing L-15 media. Media formulations without cells were used as blanks to account for media autofluorescence. All samples were allowed to grow undisturbed for 4 (Mack1) and 5 days (iBSCs), at which point wells were washed once with DPBS and frozen overnight at –80°C. Samples were then analyzed for short-term growth using direct DNA quantification with CyQUANT Cell Proliferation Assay (ThermoFisher #C7026) using manufacturer’s protocol. Emission/excitation was read at 480/520 nm on a ThermoFisher Varioskan LUX multimode microplate reader. Blank values from media-only samples were subtracted from experimental values, and these values were normalized to growth using either B8 alone (iBSCs) or 2.5 % FBS-containing L-15 media (Mack1) to compare the growth between conditions. Experiments were repeated using a narrower range of lysate concentration and using serum-free adapted iBSCs passaged 4 times consecutively in Beefy 9.

### 2.6 Adaptation and long-term growth in *Vibrio natriegens*-based media

After choosing *V. natriegens* 10 mg/mL serum-free media as a starting point for adaptation based on short-term screens, iBSCs, now at passage 60, were resuspended and seeded at 2,000 cells/cm^2^ in B8 media with 1.5 μg/cm^2^ vitronectin in a 6-well plate. After adhering for >3 h, cells were fed 2 mL B8 medium supplemented with 10 mg/mL *V. natriegens* lysate. Cells were fed *V. natriegens* lysate-based media every 2-3 days until they reached 80-90 % confluence. At this point, cells were passaged using TrypLE Express enzyme (ThermoFisher #12604013) and counted using a NucleoCounter NC-200 (Chemometec, Allerød, Denmark). Doubling time (*DT*) was calculated using **Equation 3**.

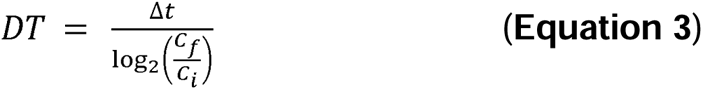

with Δ*t* being time between passages, *C_f_* being cell count at passage, and *C_i_* being total cells seeded. Cells were then centrifuged at 300 × g and seeded in a new well. This process was repeated continuously. When doubling time reached a minimum or plateaued, *V. natriegens* concentration was doubled, until doubling time was comparable to that of iBSCGM, which was when *V. natriegens* was at 40 mg/mL (VN40). At this point, VN40-adapted cells were seeded in triplicate in a 6-well plate and fed VN40. Cell doublings were measured at each passage for each replicate. This same process was repeated simultaneously for Beefy 9-adapted cells fed with Beefy 9 (as control). Cells were periodically imaged using phase contrast microscopy (KEYENCE, BZ-X700, Osaka, Japan; Olympus, CKX53, Tokyo, Japan)

### 2.7 Characterization and differentiation of VN40 adapted cells

After 20 passages in VN40 media, iBSCs were allowed to adhere in triplicate in 48-well plates at 6000 cells/cm^2^ and their satellite cell identity was characterized via staining for Paired-box 7 (Pax7). Cells were fixed with 4 % paraformaldehyde in PBS (ThermoFisher #AAJ61899AK) for 20 min, washed with DPBS containing 0.1 % Tween-20 (ThermoFisher, BP337), permeabilized for 10 min with DPBS containing 0.1 % Triton X-100 (ThermoFisher, BP151), washed again with DPBS+Tween-20, and blocked for 30 min using a blocking solution containing DPBS with 5 % goat serum (ThermoFisher #16210064) and 0.05 % sodium azide (Sigma #S2002). A primary antibody solution containing 1:500 Pax7 antibody (ThermoFisher #PA5-68506) and 1:1000 actin stain Phalloidin 488 (ThermoFisher A12379) in blocking solution was added to cells and incubated overnight at 4°C. Cells were then washed with DPBS+Tween-20 and blocked for 30 min once again. A solution containing 1:500 Pax7 secondary antibody (ThermoFisher #A11072) and 1:1000 nuclear stain DAPI (ThermoFisher #D1306) in blocking solution was added to cells and incubated for 1 h at room temperature. Cells were then washed with DPBS+Tween-20 and kept in DPBS for imaging. Imaging was carried out using fluorescence microscopy (KEYENCE, BZ-X700, Osaka, Japan). Stemness was quantified using a Celigo automated image cytometer (Revvity, Waltham, MA) to determine cells positive for Pax7. This was repeated for cells maintained in iBSCGM without VN40 adaptation to determine if adaptation altered iBSC satellite cell phenotype. Cells were differentiated by seeding at 5,000 cells/cm^2^ in 48-well plates, feeding with VN40 media, and initiating differentiation when cells appeared confluent ∼4 days after initial seeding. Differentiation media contained DMEM+Glutamax supplemented with 2 % horse serum (ThermoFisher #16050130), 0.5 mg/mL recombinant human albumin, 1× ITS-X (ThermoFisher #51400056), 0.5 mM LDN193189 (Sigma #SML0559), 1 % antibiotic/antimycotic, and 2.5 μg/mL puromycin. Differentiation continued for 18 days with cells fed fresh differentiation media every 2–3 days. After mature myotubes were observed, cells were fixed, permeabilized, and blocked as described. Cells were then stained using myosin heavy chain (MHC; Developmental studies hybridoma bank #MF-20, Iowa City, IA, USA) and 1:200 desmin (Abcam #ab15200, Cambridge, UK) primary antibodies diluted in blocking solution.

Secondary antibodies for MHC (ThermoFisher #A32723; 1:200) and desmin (ThermoFisher #A11072; 1:500) along with 1:1,000 DAPI, diluted in blocking buffer, were then applied as described. Imaging was carried out using fluorescence microscopy (KEYENCE, BZ-X700, Osaka, Japan).

### 2.8 Sample preparation for metabolomics analysis

A 100 % methanol extraction was performed on the lysate. Methanol and VN lysate were mixed in a 3:1 proportion and vortexed. The mixture was centrifuged at 15,000 × g at 4 °C for 15 min. Then, 250 μL of the supernatant was transferred to each Eppendorf tube. The samples were dried by speed vacuuming (Eppendorf Vacufuge^TM^) and stored at –20 °C. Prior to the LC-MS run, samples were reconstituted by adding methanol and water at 1:1 proportion to dried pellet, mixed by vortexing, and then centrifuged at 15,000 x g at 4 °C for 15 min (ThermoScientific Sorvall Legend X1R Centrifuge). Finally, 100 μL of the supernatant was transferred to sample tubes for LC-MS.

### 2.9 Liquid chromatography-tandem mass spectrometry (LC-MS/MS)

Untargeted analysis of metabolites present in the extracted lysate samples was preformed using information-dependent acquisition (IDA) experiments on a quadrupole time-of flight (TOF) mass spectrometer (TripleTOF 5600+, AB Sciex, Framingham, MA, USA) with an electrospray ionization (ESI) source. IDA experiments include a single TOF MS scan corresponding to four dependent product ion (MS/MS) scans based on the highest intensity unique mass. Fragmentation was triggered when precursor ion counts rose quickly over several scans to ensure ions were selected near the top of their LC peaks. A total of four methods was applied to each sample using either a hydrophilic interaction chromatography (HILIC) column (Phenomenex Luna NH2, Torrance, CA, USA) or a reverse-phase column (Phenomenex Synergi Hydro-RP, Torrance, CA, USA) in both positive and negative ionization modes. The chromatographic separation was performed with a binary pump HPLC system (1260 Infinity, Agilent, Santa Clara, CA, USA). Chromatographic gradient methods and mobile phases described in supplementary information.

### 2.10 Feature annotation

LC-MS features were annotated using the Biologically Consistent Annotation of Metabolomics Data computational tool (BioCAn)(Alden et al., 2017). Briefly, BioCAn assess potential annotations of *m/z* values and MS/MS spectra using the following annotation tools: CFM-ID(Allen, Pon, Wilson, Greiner, & Wishart, 2014), NIST17 (NIST/EPA/NIH, 2017), MONA (https://massbank.us/), and HMDB(Wishart et al., 2013) Features could be assigned multiple annotations either by a single annotation tool or by disagreement between assignments designated by different tools. To determine the most likely annotation for a feature, the feature was mapped to a metabolite in the *V. natriegens* metabolic network and assigned an annotation score by BioCAn. Annotation scores were based on agreement of the annotation assignment by the integrated databases as well as confidence in annotation of neighboring metabolites within the metabolic network.

### 2.11 Metabolomics data analysis

The peak area corresponding to the integrated area under the curve (AUC) of each ion chromatograph was blank subtracted to reduce noise. Each peak area was then normalized to the total ion chromatogram (TIC) for each replicate (n = 3). Unannotated features were subsequently removed. In addition, features showing annotation scores < 1.0 were removed due to the low confidence in the annotation. Finally, annotated features showing negative (blank subtracted) peak areas were removed as their concentration is likely lower than the limit of detection for the applied method. The remaining metabolites were then sorted by peak area.

### 2.12 Cost analysis

We compared the cost of VN lysate containing media with BSCGM and B9 media. The details of the cost comparison are in supplementary information.

## 3. RESULTS

### 3.1 Mechanical lysis of gram-negative bacteria yields extracts with high protein content

The overall process of lysate production and screening on iBSCs and the Mack1 cell line is illustrated in the schematic diagram shown in **Figure 1A**. The Gram-negative bacteria *E. coli* Nissle (“ECN”), *E. coli* BL21 (“BL21”), and *V. natriegens* (“VN”), along with the yeast strain *S. cerevisiae* CEN.PK (“CEN.PK”), demonstrate relatively higher protein yields per liter of culture compared to Gram-positive bacteria *L. lactis* (“LL”) and *L. plantarum* (“LP”) (**Figure 1B**). This discrepancy can be attributed to their thick peptidoglycan cell wall, which makes them more resistant to lysis (de Bruin et al., 2019).

We analyzed the protein profiles of lysates obtained from the microbial strains using SDS-PAGE to assess their composition. **Figure 1C** illustrates the results for the lysates from *V. natriegens*, both *E. coli* strains, *B. subtilis*, *L. plantarum*, *L. lactis*, and *S. cerevisiae*. Each lane shows distinct protein banding patterns, highlighting the differences in protein composition across species.

### 3.2 Screening helps identify compatibility between lysates and cell lines

The iBSCs used in this study were extensively characterized in our previous studies (Stout et al., 2023a; Stout et al., 2024). As an initial screen for iBSC growth in serum-free lysate-based media, cells were tested directly after passaging in serum-containing growth medium (iBSCGM). Performance (cell growth) was compared to previously established Beefy 9 serum-free medium (Stout et al., 2022), which contains similar ingredients as our tested formulations, except the recombinant human albumin present in Beefy 9 was replaced with microbial lysates. The results (**Figure 2A**) show that many of the lysates were inhibitory to cell growth after 5 days of exposure. All lysates elicited decreased cell growth at higher concentration (up to 500 µg/mL). Lysates from VN (10 µg/mL) and LP (10 and 100 µg/mL) exhibited short-term cell growth similar to that of the Beefy 9 control.

**Figure 2.**
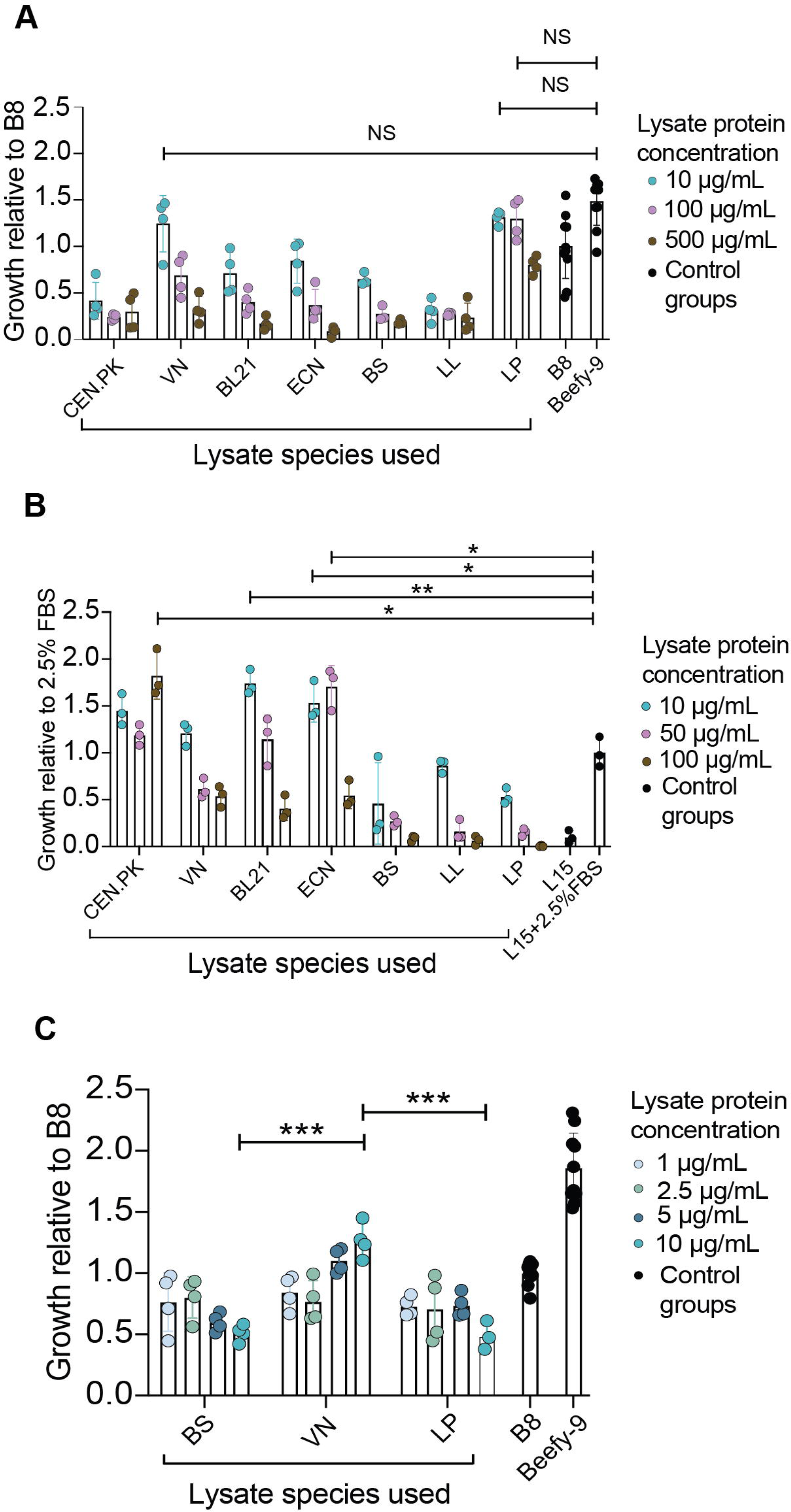
Short-term proliferation assay of iBSCs and Mack1 cells using various microbial lysates. **A)** iBSC (not serum-free adapted) proliferation screen (5-day incubation) on microbial lysate-based serum-free media shows VN and LP lysates performing similarly to Beefy 9 medium (positive control). B8 medium (Beefy 9 without rAlbumin) was used as negative control. Lysate-based media created through addition of respective lysates to B8 media. Data is from CyQuant DNA staining kit. n=4 (experimental groups) or 10 (controls). **B)** Mack1 (not serum-free adapted) proliferation screen using an L-15 medium supplemented with 2.5 % FBS (negative control) and additional microbial lysates. n=3. CEN.PK was well tolerated at different concentrations. **C)** iBSC (4× serum-free Beefy 9 passages) proliferation screen using selected lysates at lower concentrations. n=4 (experimental groups) or 10 (controls). Lysate formulations were compared to one another to determine candidates for long-term growth studies, showing VN as top contender. All presented data is mean values ± SD. Statistical analysis by unpaired t-test with Welch’s correction in which p > 0.05 (NS), p < 0.05 (*), 0.01 (**), and 0.001(***).

We also screened the short-term growth-promoting effects of lysate-based reduced-serum (2.5% FBS) media on Mack1 cells (**Figure 2B**). Given that all lysates at 500 µg/mL showed detrimental effects on iBSC growth, we tested a lower range of lysate protein concentrations for Mack1 cells. Unlike with iBSCs, VN at concentrations above 10 μg/mL and LP at all concentrations inhibited short-term Mack1 cell growth. However, CEN.PK, BL21, and ECN lysates significantly improved the short-term growth of the cells. Specifically, Mack1 cells performed best in the presence of CEN.PK (up to 100 µg/mL), BL21 (10 µg/mL), and ECN (10 µg/mL and 50 μg/mL) lysates. At 100 µg/mL, CEN.PK lysate nearly doubled short-term growth of Mack1 cells, having a similar effect as 7.5% FBS-containing media as shown by Lim et al. (2024), suggesting this lysate can replace 5% FBS in media without detrimental effect.

Despite our success with Mack1 cells, we found greater success in adaptation of iBSCs to serum-free media using lysates and proceeded with subsequent studies using only iBSCs. Since serum-free adaptation significantly alters cellular phenotype and gene expression (J.Y. Lee et al., 2022), transferring cells directly from serum-containing media to lysates may not be an appropriate strategy to assess the long-term potential of lysates. So, next we re-screened the lysates using iBSCs grown in Beefy 9 (serum-free) medium (4 passages) with select lysates (**Figure 2C**). The bovine cells were grown in lysate concentrations lower than those of the first screen, since concentrations above 10 µg/mL were shown to be cytotoxic. The results showed that cells cultured in serum-free conditions responded differently to lysates than those cultured only in serum-containing media. Cells cultured in VN media at 10 µg/mL outperformed those grown at this concentration of BS and LP. Cell growth improved with increased VN lysate concentration between 2.5 and 10 µg/mL, while the opposite effect was seen in cells grown in LP and BS. While initial screens found that non-adapted cells proliferated in LP media, these results did not translate to cells adapted to serum-free media. Given that VN performed well with serum adapted and un-adapted cells, we chose it for further investigations.

### 3.3 iBSCs can adapt to long-term growth on *V. natriegens* lysate containing media

iBSCs were cultured directly from BSCGM into serum-free B8 medium supplemented with 10 µg/mL VN lysate. Growth was monitored throughout several passages and lysate concentration in feeding media was increased when cell growth slowed. This strategy led to increases in cell growth rate and was used to slowly adapt cells to a medium with VN lysate protein concentration of 40 µg/mL (**Figure 3A**), dubbed “VN40”. At this concentration, cell growth rate plateaued, and doubling time reached 24.9 h, similar to those observed using serum-containing BSCGM (21.9 ± 3.1 h) during the course of the study. Overall growth kinetics of iBSCs grown in VN40 over 25 days were similar to those grown in 20% FBS BSCGM medium – ∼19 total cell doublings in VN40 compared to ∼22 in BSCGM (Stout et al., 2022). This is a significant increase from the ∼14 doublings shown in Beefy-9 and implies VN lysate can completely replace FBS in iBSC growth media. Cells showed an increase in viability between 10 µg/mL VN feeding (avg. 71.5 % viability) versus 40 µg/mL feeding (avg. 93.9 % viability), though this may have been a factor of initial population enrichment during adaptation to lysate-based medium. In all, the process of adapting cells to VN40 took approximately 2.5 months (passage 1 to passage 14), in line with previous durations of adapting mammalian cells to serum-free media (Hartman et al., 2007; Hua et al., 2022; Miki & Takagi, 2015). Adaptation of iBSCs to VN40 was expedited by passaging cells directly into 20 µg/mL VN medium (**Figure 3B**), reducing total adaptation time to 4 weeks. After adaptation of cells in VN40 medium, cells were grown in triplicate to compare to growth in serum-free Beefy 9 media. Both cell populations were passaged continuously over 24 days with collective cell doublings recorded at each passage (**Figure 3C**). Cells grown in VN40 significantly outgrew cells grown in Beefy 9, with higher collective cell doublings and lower average doubling times (31.2 ± 4.8 h for VN40; 43.1 ± 8.7 h for Beefy 9). An increase of VN lysate content in the media beyond 40 µg/mL hindered cell growth compared to VN40 (**Figure S1**). Cells had higher confluence after 3 days of growth in VN40 compared to Beefy 9 (**Figure 3D-E**). VN40 also resulted in less observable lipid accumulation compared to Beefy 9 (**Figure 3F-G**). Beefy 9 has been shown to cause aberrant lipid accumulation (Stout et al., 2022), potentially due to induced insulin resistance from the high concentration of insulin in B8, or via adipogenic signaling. Components of VN40 may be preventing such insulin resistance and/or adipogenic signaling from taking place, which could be beneficial in long-term maintenance of myogenicty and proliferative capacity. Cells in VN40 have maintained proliferation over 108 passages and can survive long-term storage and thawing from liquid nitrogen.

**Figure 3.**
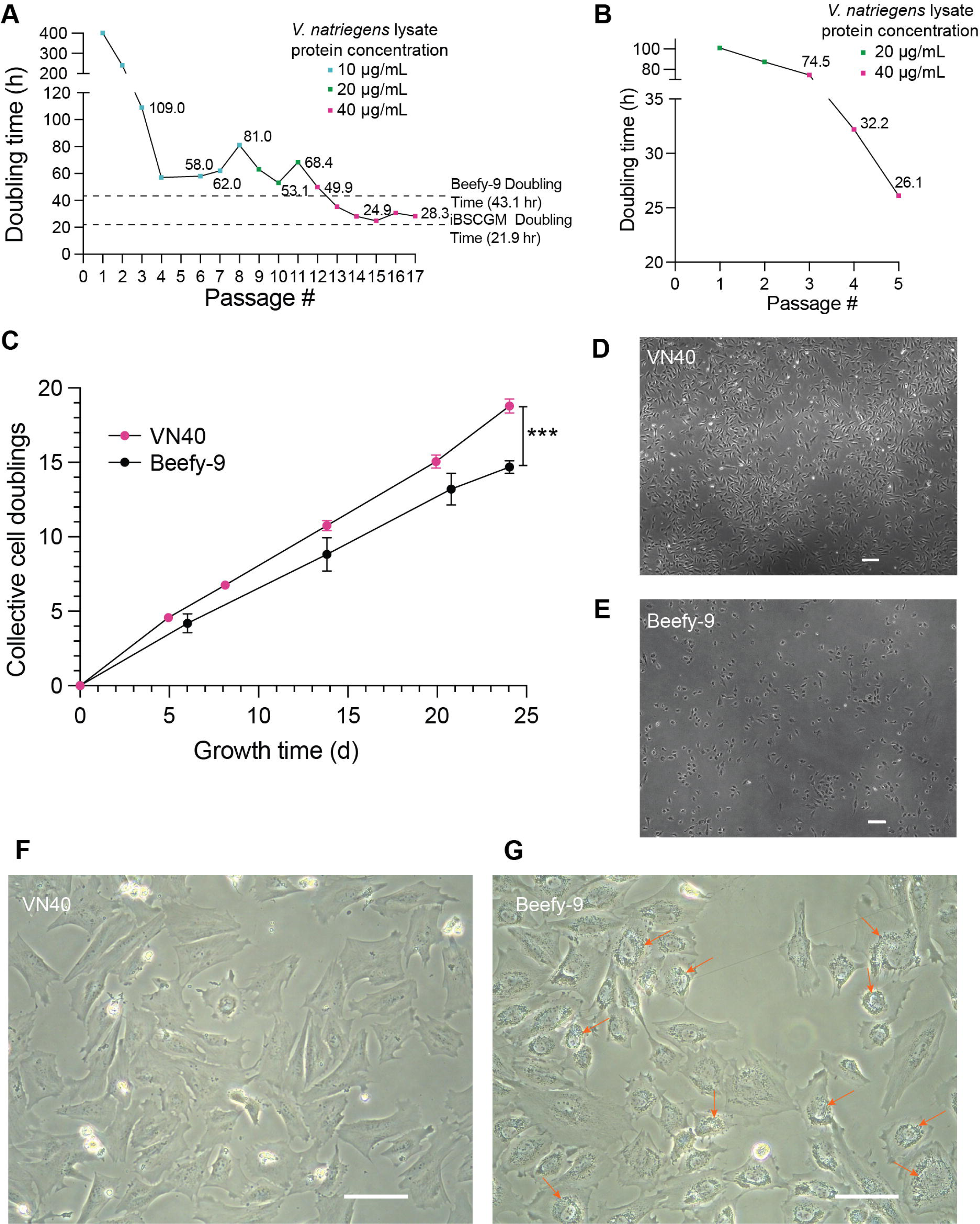
Adaptation and long-term growth of iBSCs in VN40. **A)** iBSCs grown over multiple passages in B8 medium containing increasingly concentrated *V. natriegens* lysate (corresponding to 10, 20, 40 µg/mL protein concentrations). The cells were grown in T-25 flasks and passaged at 80– 90 % confluence. Dotted line representing doubling times in Beefy 9 & BSCGM (General Media for iBSCs containing 20 % FBS) collected throughout this study. **B)** iBSC adaptation to *V. natriegens* lysate-containing media beginning with 20 µg/mL protein concentration. **C)** iBSCs grown continuously in VN40 and Beefy 9 media with cells counted at each passage to determine cumulative cell doublings. Cells grown in triplicates in 6 well plates for each condition. Significance indicated for final cumulative cell doublings. All presented data is mean values ± SD. Statistical analysis by unpaired t-test with Welch’s correction in which p < 0.001 (***). **D-E)** Cells after the final experimental passage (passage 82 overall, passage 22 in serum-free media) grown for 3 days and imaged by phase contrast microscopy, showing improved growth of VN40 (**D**) compared to Beefy 9 (**E**). Scale bar = 200 µm. **F-G)** Magnification increased to show additional detail, with orange arrows indicating increased cytoplasmic lipid accumulation in cells grown in Beefy 9 (**G**) compared to VN40 (**F**). Cells were quantified manually or using an automated cell counter. Scale bar = 100 µm.

### 3.4 High and low molecular weight components of *V. natriegens* lysate are additive in promoting iBSC proliferation

To determine if a specific molecular weight fraction contributed to the mitogenic effect of VN lysate-based media, we fractionated the lysate by ultrafiltration using a 3 kDa molecular weight cutoff membrane. iBSC growth was determined via the same protocols as earlier, with short term growth compared to that in B8 (negative control) and Beefy 9 (positive control). Formulations using the protein-rich high molecular weight (HMW) >3 kDa fraction, low molecular weight (LMW) <3 kDa fraction, and whole-cell VN lysate, were prepared at various concentrations in B8 media. For total lysate and HMW fractions, protein concentrations were determined using Bradford assay, as described previously. Since the protein component of the LMW fraction was below the detection limit for Bradford assay, we used the same composition as would have been required for the whole cell lysate. The results (**Figure 4A**) showed that for concentrations of VN protein equal to or above 40 µg/mL, whole cell lysates out-performed HMW or LMW fractions alone. It can be concluded from this data that no individual fraction of VN lysate was cytotoxic compared to whole-cell lysates, and that larger proteins act together with small molecule metabolites to have an overall pro-mitogenic effect on iBSCs. This has been shown in prior work in which metabolomics and proteomics showed the indispensability of both amino acid and protein fractions in *Auxenochlorella pyrenoidosa* extracts for goldfish muscle cell growth (Dong et al., 2023). Whole-cell lysates of VN40 outperformed individual fractions, suggesting that the proteins and metabolite fractions have an additive impact (**Figure 4B**). Ultimately, our data shows the importance of using unfractionated VN40 in stimulating iBSC growth.

**Figure 4.**
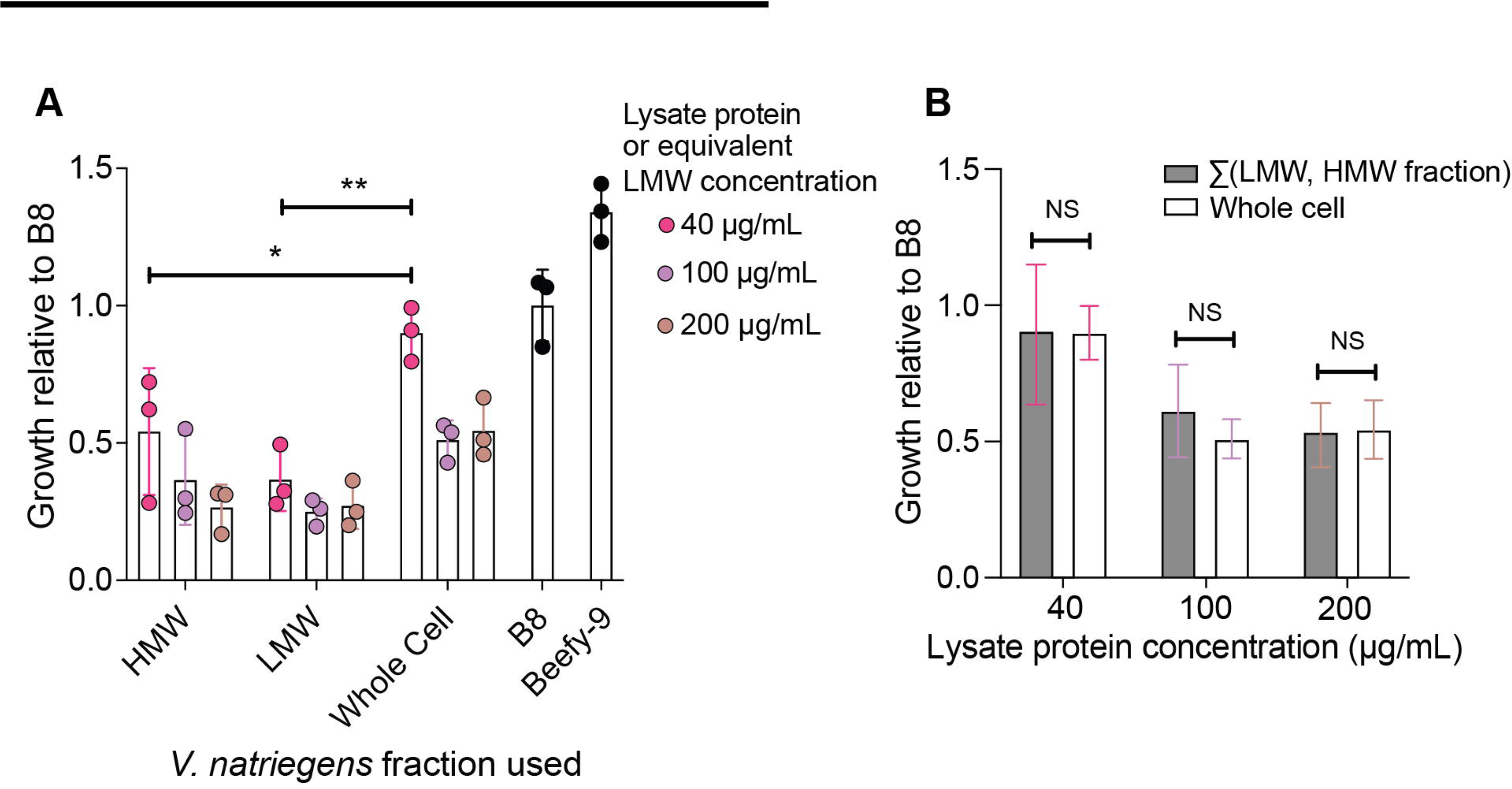
Assessing the role of HMW and LMW *Vibrio natriegens* lysate fractions on iBSC growth. **A)** The growth of iBSCs using a short-term screen (5 days) demonstrates that whole-cell lysates support growth more effectively than either the >3 kDa (HMW) or <3 kDa (LMW) fractions alone. HMW and LMW samples represent retentate and filtrate, respectively, that were added to B8 media at protein concentrations indicated (<3kDa concentration assessed by adding low molecular weight fraction representative of corresponding whole cell lysate). Beefy-9 media used as positive control, B8 media used as negative control. Data is from CyQuant DNA staining kit. **B)** iBSC proliferation in whole-cell VN40 medium compared to the sum of HMW and LMW growth indicates additivity of their relative contributions to iBSC growth. All presented data is mean values ± SD. Statistical analysis by unpaired t-test with Welch’s correction in which p > 0.05 (NS), p < 0.01 (**).

### 3.5 VN40 cultured iBSC maintain stemness and can be differentiated

We validated continued satellite cell phenotype via Pax7 expression (Relaix et al., 2005) of iBSCs after 22 passages in VN40, showing that expression was maintained throughout serum-free adaptation (**Figure 5A-B**). According to quantitative imaging cytometry, cells were 99.7% positive for Pax7 after adaptation (**Figure 5C**), indicating maintenance of a pure satellite cell population. After validation of phenotype, we successfully differentiated VN-40 adapted cells into mature multi-nucleated myotubes expressing both Desmin and Myosin Heavy Chain proteins (**Figures 5D-K**), indicative of early-stage and late-stage differentiation respectively (Capetanaki et al., 2022). This implies that VN40-adapted cells maintain their myogenicity after extensive serum-free passaging, essential for creating functional cultivated meat products (M. Lee et al., 2024). Cells differentiated after slow, gradual adaptation (10 µg/mL → 20 µg/mL → 40 µg/mL VN lysate) (**Figure 5D-G**) differentiated more slowly and less robustly than cells differentiated after a quicker adaptation (20 µg/mL → 40 µg/mL) (**Figure 5H-K**). This is likely due to the impact of higher passage number on myogenicity (Park et al., 2023). Differentiation media was optimized to contain ITS-X (Messmer et al., 2022) and LDN-193189 (Horbelt et al., 2015), which were necessary for differentiating these overall high passage (>p70) cells. Ultimately, we have arrived at a bacterial lysate-based serum-free formulation which allows for successful proliferation and differentiation of iBSCs.

**Figure 5.**
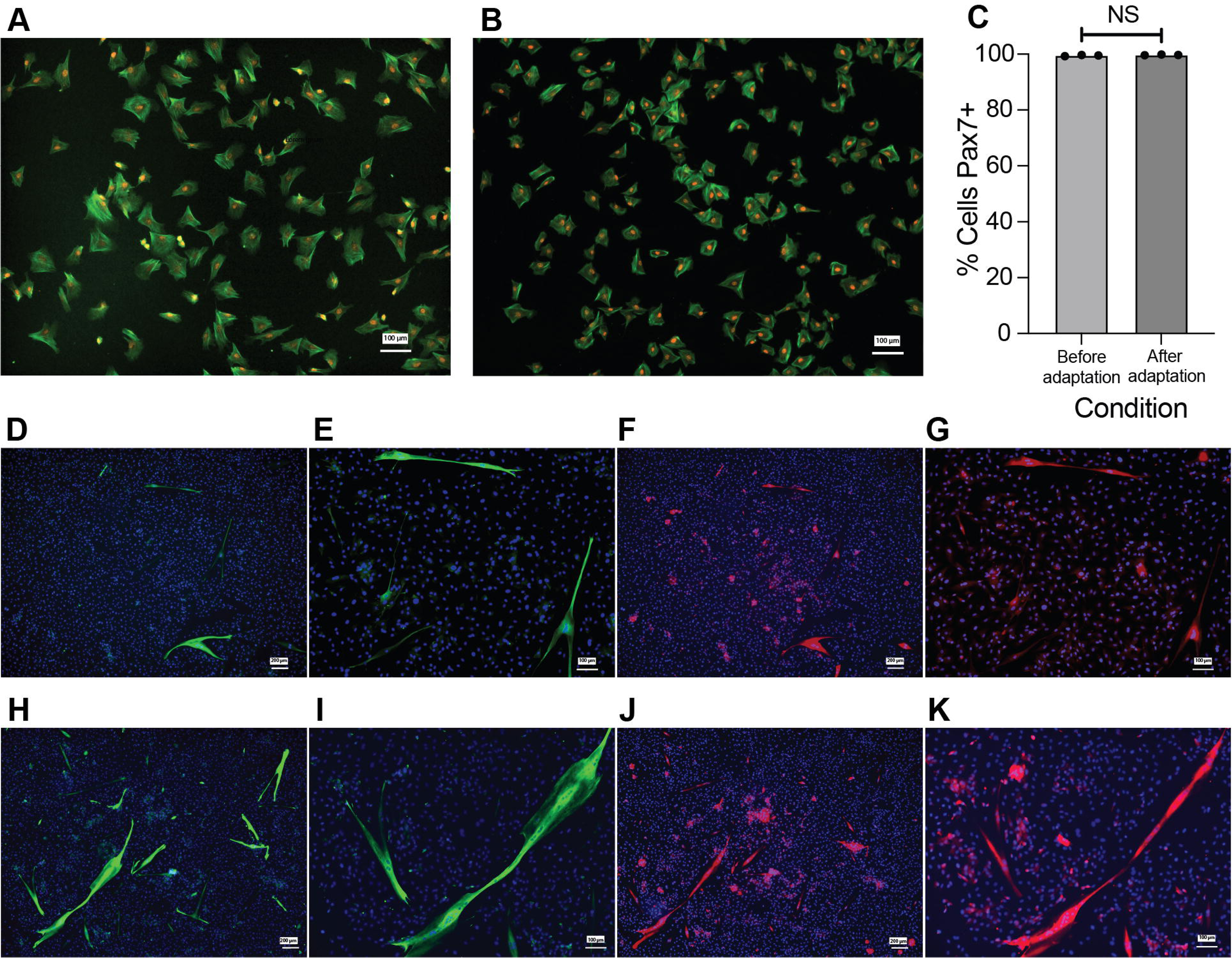
Stemness and differentiation of VN40-adapted iBSCs. **A-B)** Immunofluorescent staining of iBSCs before (**A**) and after (**B**) adaptation and serial passaging (22 passages) in VN40 medium show no change in satellite cell phenotype after adaptation. Cells were stained with phalloidin (green) to indicate actin expression and anti-Pax7 (orange). Scale bar = 100 µm. **C)** Quantification of Pax 7+ cells before and after adaptation and serial passaging (22 passages) in VN40 medium. % Cells Pax7+ determined by (number of Pax7+ cells)/(number of actin+ cells). Statistical analysis by unpaired t-test with Welch’s correction in which p>0.05 (NS). **D-G)** Immunofluorescent staining of gradually adapted iBSCs (p98) after 18 days differentiation in differentiation media (2 % horse serum, 50 µg/mL rAlbumin, 0.5 µM LDN-193189, ITS-X). **H-K)** Immunofluorescent staining of rapidly adapted iBSCs (p70) after 13 days differentiation in differentiation media. Results show successful differentiation of cells expressing early-stage (Desmin, green) and late-stage (myosin heavy chain, red) differentiation markers, with more robust differentiation of rapidly adapted cells. Cells were stained with anti-desmin, MF20 (for myosin heavy chain), and DAPI (blue) to stain nuclei. **D,F,H,J:** low magnification, Scale bar = 200 µm; **E,G,I,K:** high magnification, Scale bar = 100 µm.

### 3.6 LC-MS/MS identifies major metabolites in *V. natriegens* lysate

VN lysate was analyzed for metabolite composition using a series of untargeted LC-MS/MS assays. The untargeted analysis detected a total of 11,306 features using a combination of HILIC and reverse phase chromatography in both positive and negative ionization modes (four total methods) The number of detected features in our study aligns with typical results from untargeted metabolomic analyses. This is due to the generation of degenerate features, such as those resulting from adduct formation, dimerization, isotopic variants, and fragmentation variation, which can inflate the total number of features derived from single unique metabolites by up to 90% (Mahieu et al., 2017). A previously developed automated tool (BioCAn) was used to annotate the features, resulting in a total of 403 metabolite annotations that mapped to known metabolites in VN. Similar network-based feature annotation tools demonstrate comparable identification capacities to BioCAn. For instance, MetDNA utilizes a reaction network-based recursive algorithm to sequentially identify features utilizing a surrogate MS/MS spectra from an initial set of seed metabolites. In a study on *Drosophila* samples, MetDNA annotated 1,314 metabolites out of 18,320 detected features with an accuracy of 77.5% (Shen et al., 2019). A significant limitation of network-based annotation algorithms is the dependence on the understanding of the organism’s metabolic network. As a result, model organisms with more comprehensive and curated genomic information and metabolic function tend to achieve greater metabolite coverage, while lesser studied organisms, such as *V. natriegens,* may face more limited metabolite annotations. Although BioCAn uses biological network data to reduce false positives, errors may still occur. Thus, only annotations with a score greater than 1.0 were further considered. An annotation score >1.0 indicates strong agreement across integrated databases, as well as similar agreement in the annotation of metabolites within two reactions of the annotation in the metabolic network. A total of 155 metabolites were identified across the 4 methods with annotation scores ≥1.0 (**Table 2**), which included 80 unique metabolites. An analysis of commonality between top metabolites detected by each method can be seen in **Figure S2.** Interestingly, purine and pyrimidine derivates and aromatic amino acids show the greatest area under the curve, suggesting potentially elevated concentrations of these metabolites in VN lysate extracts.

**Table 2.**
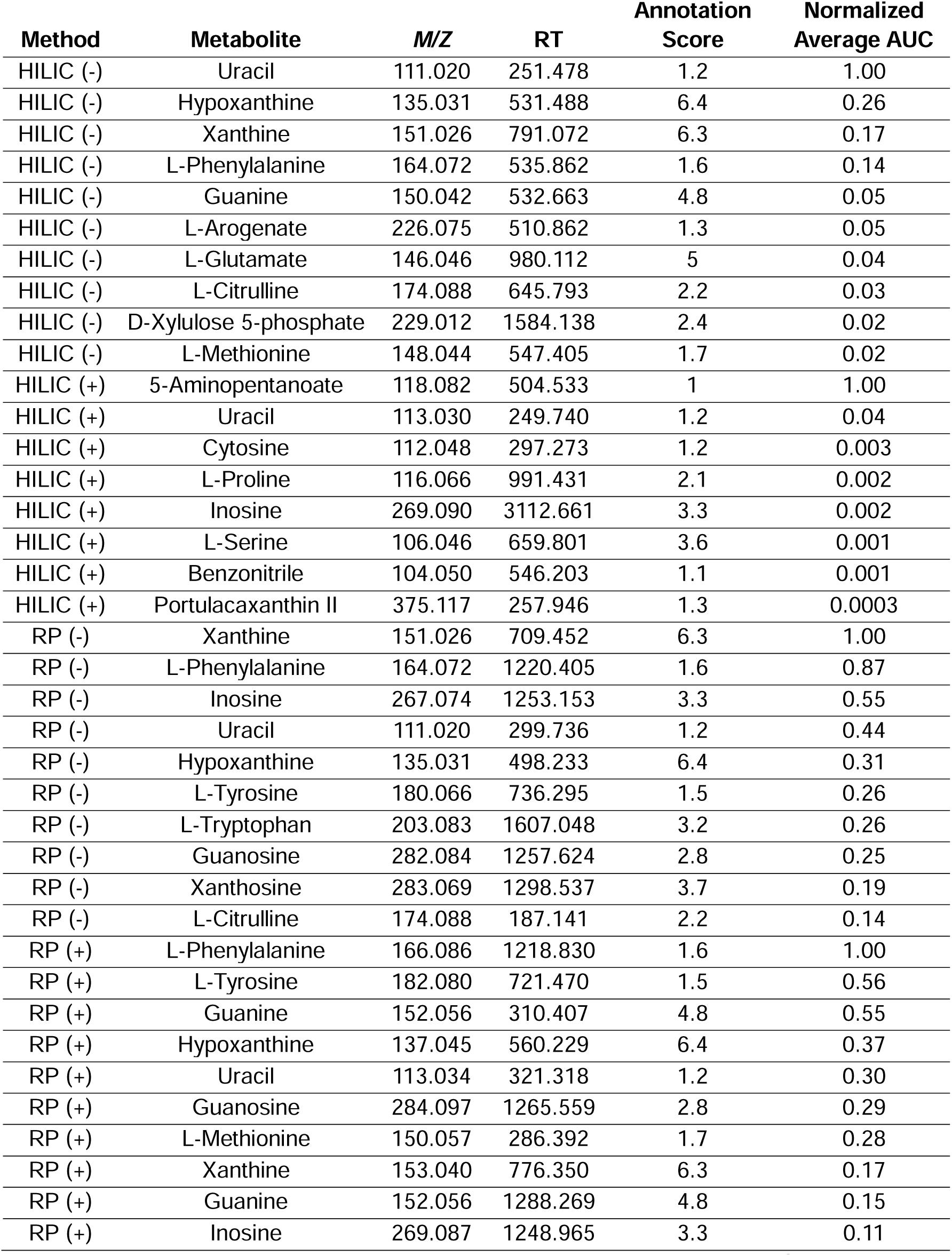
Annotated metabolites showing greatest AUC detected in each LC-MS/MS method. Each metabolite’s area is normalized to that with the highest AUC for each LC-MS method (n=3). Top 10 greatest AUC metabolites shown for each method, except HILIC (+) which found only 8 metabolites with annotation score >1.0.

### 3.7 VN40 component costs at lab scale are competitive relative with Beefy 9

Overall production cost of cultivated meat includes at least three main factors – bioreactors, labor, and cell culture medium. For this study, we have focused on the economics of FBS, a significant factor in cell culture medium content and costs. Understanding the cost of VN lysate compared to FBS is necessary to predict the sustainability of cultivated meat made using lysates as key media components. VN lysate at 40 μg/mL used as a replacement of FBS is sustainable and cost effective at $2 per liter compared to $264 of FBS per liter, a 99.2 % price reduction, at lab scale (**Table S2 & S3**). Thus, based on the calculations, VN40 reduces media cost from $323.30 to $158.60, a 51% reduction (**Figure 6**). This accounts for only reagent costs involved in the production of *V. natriegens* lysate in a lab. On an industrial scale, while the relative advantage of VN lysate over FBS may remain consistent, the absolute cost of mass-producing VN lysate should be lower than that calculated here.

**Figure 6.**
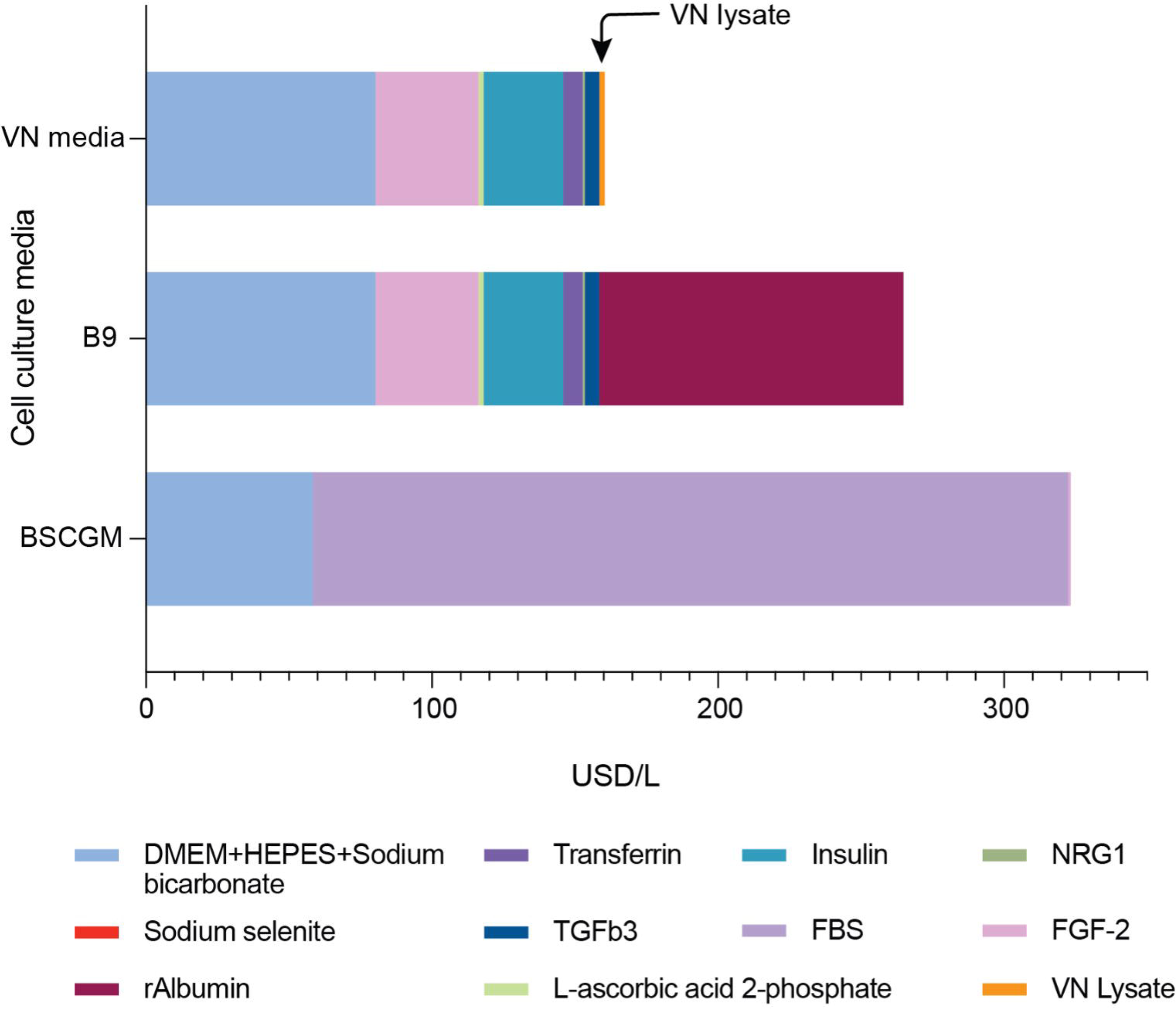
VN lysate is less expensive at lab scale than FBS, making VN40 media more economically sustainable than BSCGM. Media costs for BSC-GM, Beefy-9 and VN40 were assessed for reagents purchased at USD/L. VN media is the most economical followed by B9 and BSC-GM. FBS in BSCGM and rAlbumin in B9 are major cost-drivers for respective media compared to negligible cost of VN lysate in VN media.

## 4. DISCUSSION

Issues of increasing global food insecurity and climate crisis (Boyd et al., 2020; Hayek, 2022; Poore & Nemecek, 2018), coupled with the growing demand for meat (OECD et al., 2021), create an urgent need for more sustainable approaches to meat production. Cultivated meat, the process of growing meat *in vitro* without the need to raise livestock, is a possible alternative approach (Sinke et al., 2023; Treich, 2021). However, production of cultivated meat needs to be inexpensive, reliable, and relatively simple to compete with the low costs of traditional meat products from industrial farming.

Moreover, the reliance on FBS for cell culture is a significant roadblock to scaling and commercializing cultivated meat due to FBS’s animal origin, high cost, lot-to-lot variability, and potential as a source of contamination (Urzì et al., 2022; Zheng et al., 2006). Recombinant proteins (Kolkmann et al., 2022; Skrivergaard et al., 2023; Stout et al., 2023a; Venkatesan et al., 2022) or hydrolysates from organisms (Andreassen et al., 2020; Okamoto et al., 2022; Stout et al., 2023b; Teng, Lee, & Chen, 2023) are costly, labor intensive, and at times, still reliant on animal products or serum. Microbial extracts can be a protein-rich, inexpensive, and scalable alternatives to generating FBS-replacing media supplements. Microbial extracts have been utilized for cultivated meat production (Celebi-Birand et al., 2023; Ghosh et al., 2023; Lei et al., 2023), but still depended on serum (Lei et al., 2023), toxic bacteria (Ghosh et al., 2023), or otherwise animal-derived microbes (Celebi-Birand et al., 2023) to stimulate growth. Further, manufacturing these microbe-based serum-free media (SFM) formulations is limited by relatively slow growth rates of the chosen microbes.

In screening microbial lysates for cell line growth, we anticipated positive outcomes in either Gram positive bacteria (lacking lipopolysaccharides), or food-associated/probiotic microbes such as *S. cerevisiae* and *Lactococcus lactis.* Growth of Mack1 cells in 2.5% FBS medium supplemented with *S. cerevisiae* lysate 100 µg/mL was equivalent to that shown in 7.5% FBS-containing media by Lim et al. (2024). This means the yeast lysate can replace 5% FBS in medium without detrimental effects. The beneficial impact of *V. natriegens* lysate on iBSCs was a surprise, highlighting the importance of individualized screens for separate cell lines, potentially due to unique nutritional and signaling requirements (O’Neill et al., 2022). After screening cell growth using lysates from seven microbial strains, *V. natriegens* (a fast-growing marine bacterium) was chosen for use in long-term adaptation of iBSCs to SFM. By sequentially increasing *V. natriegens* lysate concentration in SFM, we were able to achieve maximal doubling rates of iBSCs using the VN40 formulation. This led to cell growth rates exceeding previously established Beefy 9 medium (Stout et al., 2022), while preserving iBSC phenotype and myogenicity. VN40 was able to stimulate long-term growth in iBSCs at up to 24.9 h doubling times, rivaling that of serum-containing media. The lysates were simple to produce and highly potent, stimulating cell growth at a mere 40 µg/mL. Further results indicated the importance of both low- and high-molecular weight components in the microbial extract, suggesting there is no need for process-intensive ultrafiltration of lysate fractions to achieve optimal performance as in other studies (Ho et al., 2021; Stout et al., 2023b; Teng et al., 2023). LC-MS analysis of VN lysate revealed the presence of aromatic amino acids, purine derivatives and nucleoside intermediates which are also present in FBS (D. Y. Lee et al., 2022). These compounds aid in the proliferation of iBSCs through known mechanisms. Aromatic amino acids are synthesized in cow’s rumen by fermentation (Khan et al., 2002), and can be found in FBS and basal media used for mammalian cell culture (Liu et al., 2023). The aromatic amino acids found in VN lysate could serve as nutrients for iBSCs and aid protein synthesis, cellular signaling, and/or oxidative stress response (Liu et al., 2023). The three aromatic amino acids found in VN lysate (tyrosine, phenylalanine, and tryptophan) are specifically known to be ubiquitously involved in protein synthesis, as well as synthesis of a host of metabolites vital for cell growth (Parthasarathy et al., 2018). They are also important for stimulating cell proliferation via the MAPK pathway (El Refaey et al., 2015), a pathway similarly stimulated by growth factors such as FGF, EGF (epidermal growth factor) and VEGF (vascular endothelial growth factor) (Rodrigues et al., 2010) found in FBS. Purines, pyrimidines, and their derivatives could contribute to the growth of iBSC in absence of any serum. These nucleotides found in the VN lysate, such as uracil, guanine, and cytosine, supplement precursors for DNA and RNA via the nucleotide salvage pathway in serum-free conditions lacking these metabolites (Morrison et al., 2019).

Supplementation of the same nucleosides found in VN lysate (eg. hypoxanthine, inosine, and guanosine) have shown to improve growth rates in CHO cell cultures (Morrison et al., 2019). Inosine specifically can serve as a carbon source in media lacking the nutrients of FBS (Wang et al., 2020). While our analysis did uncover several compounds with pro-mitogenic properties in the absence of serum, any imbalance of these compounds can cause cytotoxicity. This explains why a high concentration of lysates may have inhibited cell growth. An imbalance of nucleotides can cause cell growth inhibition (Diehl et al., 2022). Overabundance of the aromatic amino acids found in VN lysate can lead to cell cycle arrest (Xavier et al., 2024). Numerous other compounds in VN lysate may have become cytotoxic to cells at higher concentrations. Further optimization of individual compounds in lysate-based media is worth investigation.

*V. natriegens* is a promising ingredient in cultivated meat media, as it is the fastest growing non-pathogenic bacteria known, with a generation time of < 10 min (H. H. Lee et al., 2019). Further, its nutritional needs are both minimal and flexible (Ellis et al., 2019). Our work shows its potential to supply the cultivated meat industry with an inexpensive on-demand serum-free supplement in a potent form. Previous environmental impact assessments using microbial lysates assumed a need for microbial protein on the scale of milligrams per milliliter of media (Tuomisto & de Mattos, 2011). Our media, only requiring 40 µg/mL protein, would reduce such needs by a factor of one hundred. This would result in significant decreases in water usage, energy usage, and greenhouse gas emissions associated with cultivated meat production, which largely stem from production and processing of media ingredients (Sinke et al., 2023). This high potency would also significantly reduce cost of cultivated meat production, which are largely driven by media and additional growth factor costs (Specht, 2020).

## AUTHOR CONTRIBUTIONS

James Dolgin, Damayanti Chakravarty – Conceptualization; formal analysis; investigation; methodology; data visualization; writing & editing. Yiming Cai, Taehwan Lim, Pomaikaimaikalani Yamaguchi – Formal analysis; investigation; methodology; data visualization; writing. Sean F. Sullivan, Joseph E. Balkan, Licheng Xu, Aaron D. Olawoyin – investigation; Kyongbum Lee – funding acquisition; supervision; writing— review. David L. Kaplan and Nikhil U. Nair – Conceptualization; funding acquisition; supervision; writing – reviewing & editing.

## ACKNOWLEDGMENTS

This work was supported by funds from Tufts University Center for Cellular Agriculture Industry Consortium (to N.U.N., D.L.K., K.L.) and the USDA (2021-05678) (to D.L.K.).

## Supplemental Information

**Table S1.**
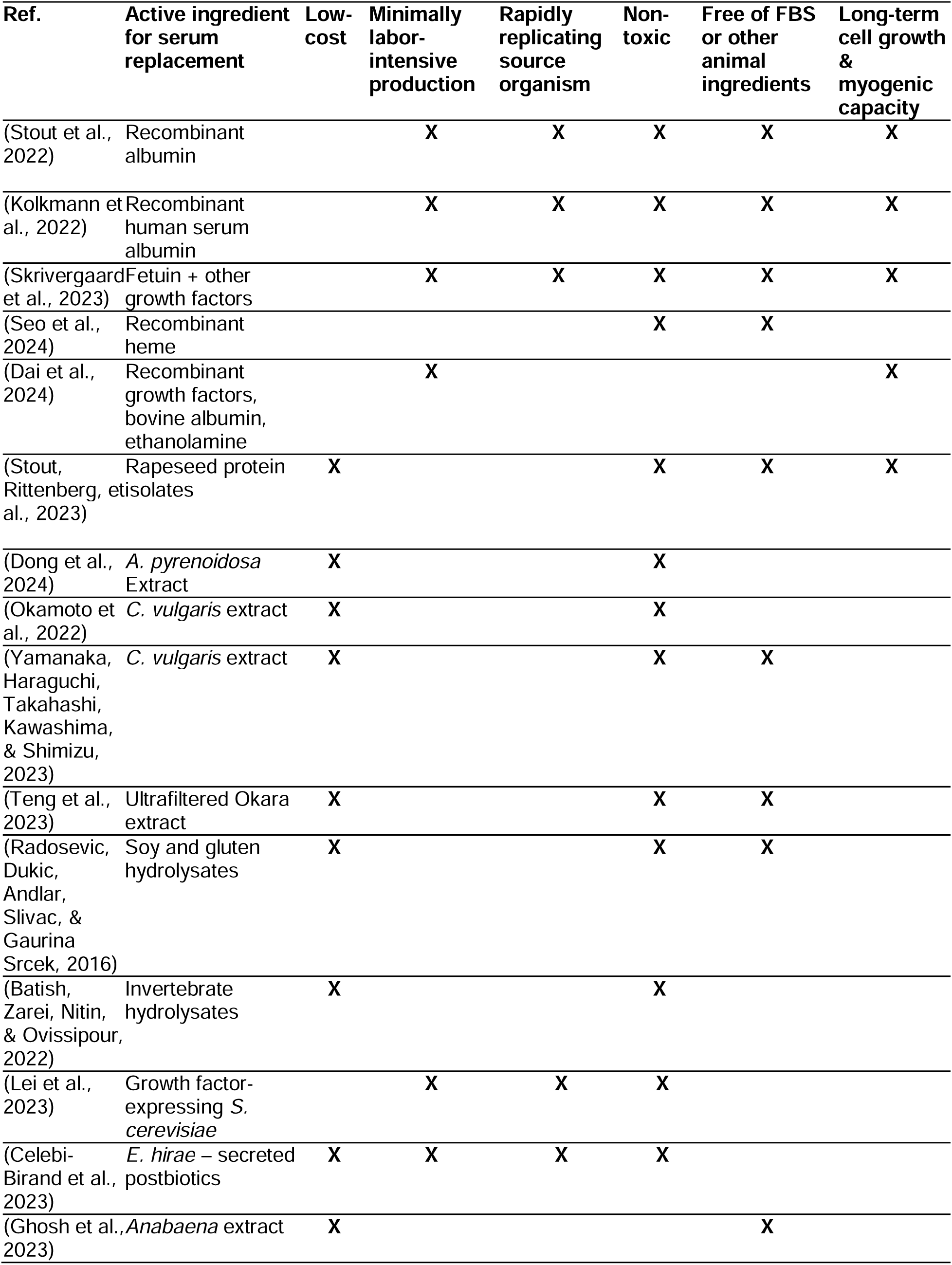
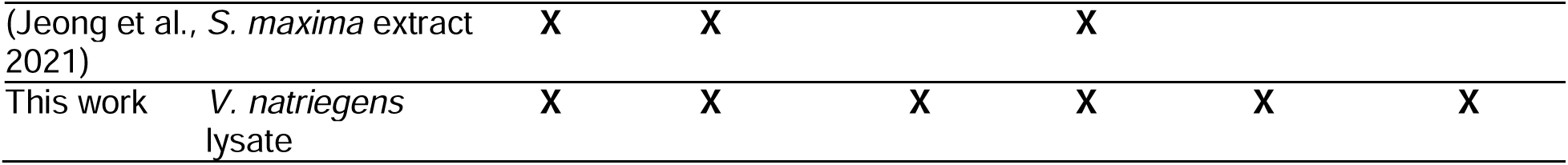
Previous research exploring cultivated meat media, with advantages of current work indicated.

**Figure S1.**
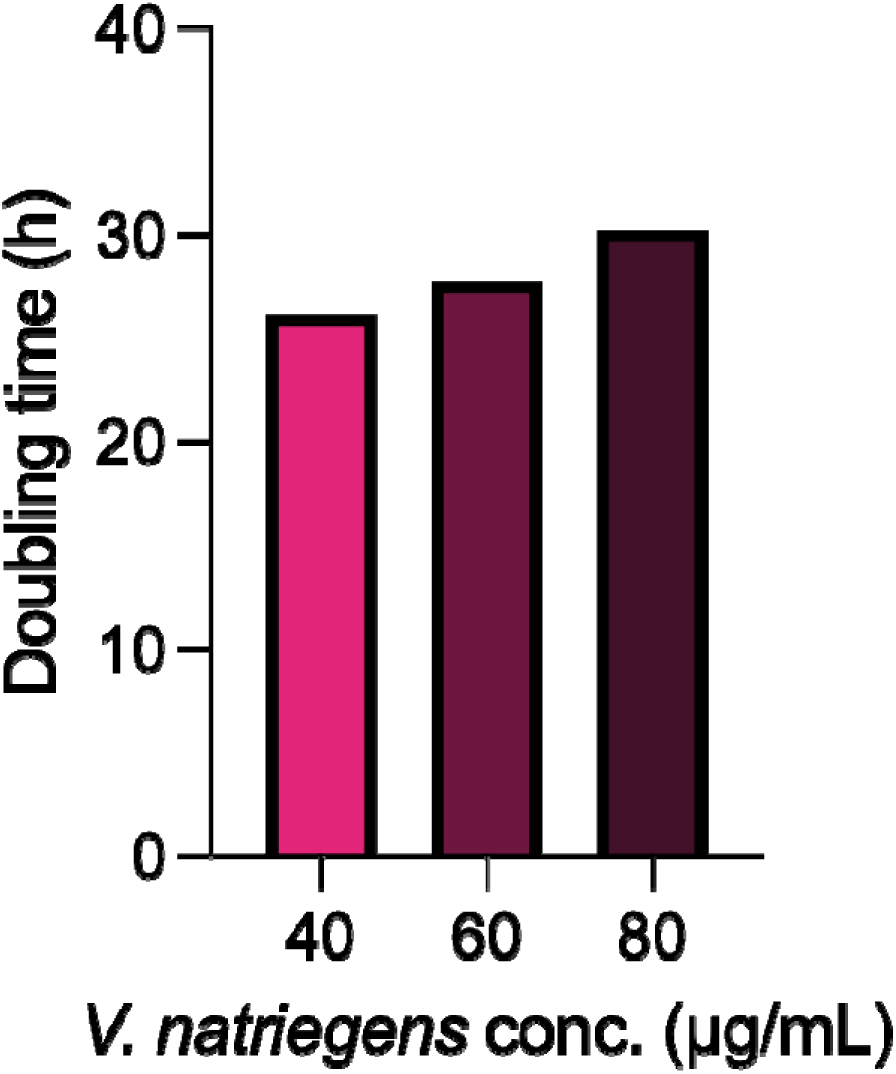
iBSCs fed serum-free media with *V. natriegens* lysate at 40 (VN40), 60, and 80 µg/mL, and resultant doubling times.

**Figure S2.**
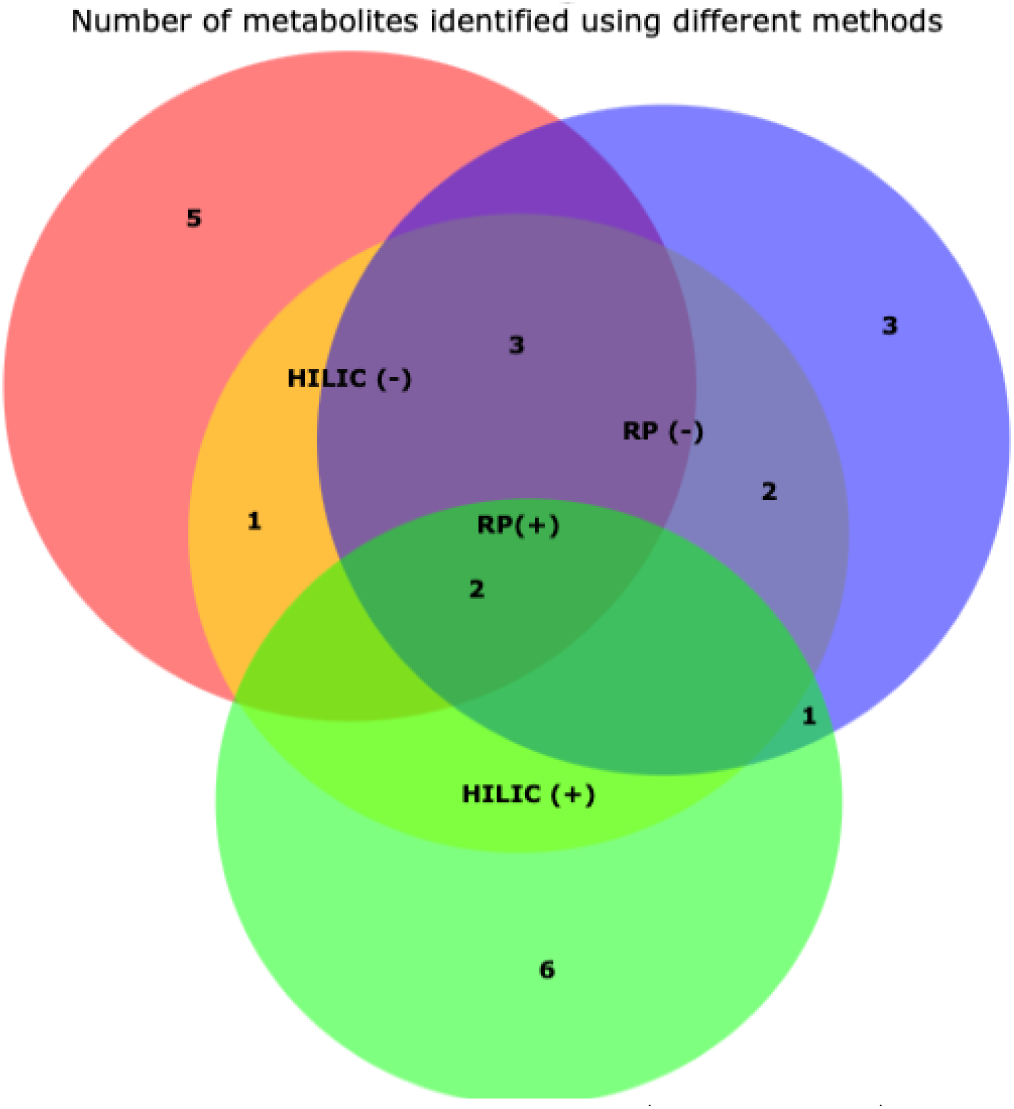
Venn diagram indicating the top 10 highest AUC (8 for HILIC+) metabolites detected by each method (HILIC (-), HILIC (+), RP (+) and RP (-) and the common metabolites among the method groups.

**Table S2.**
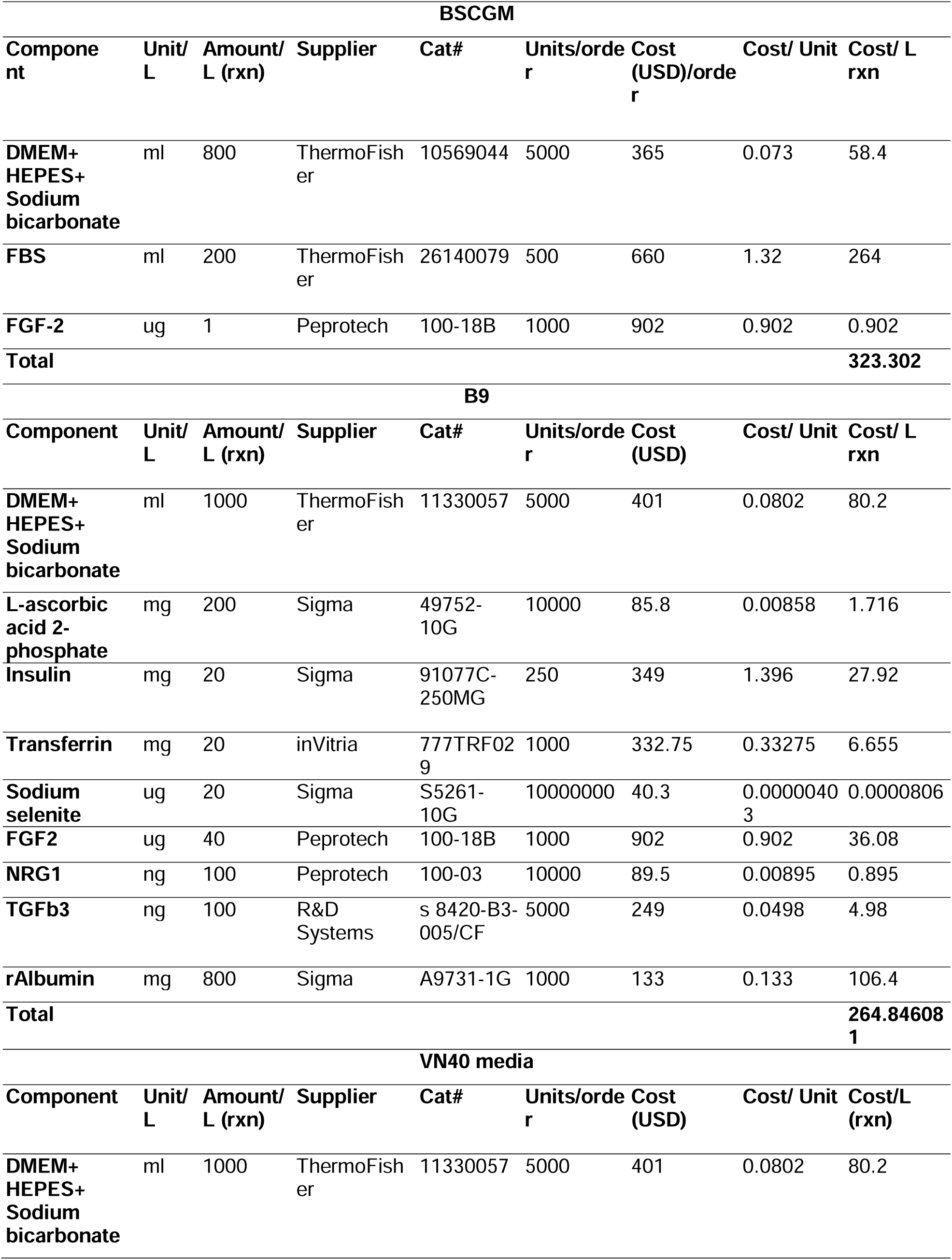
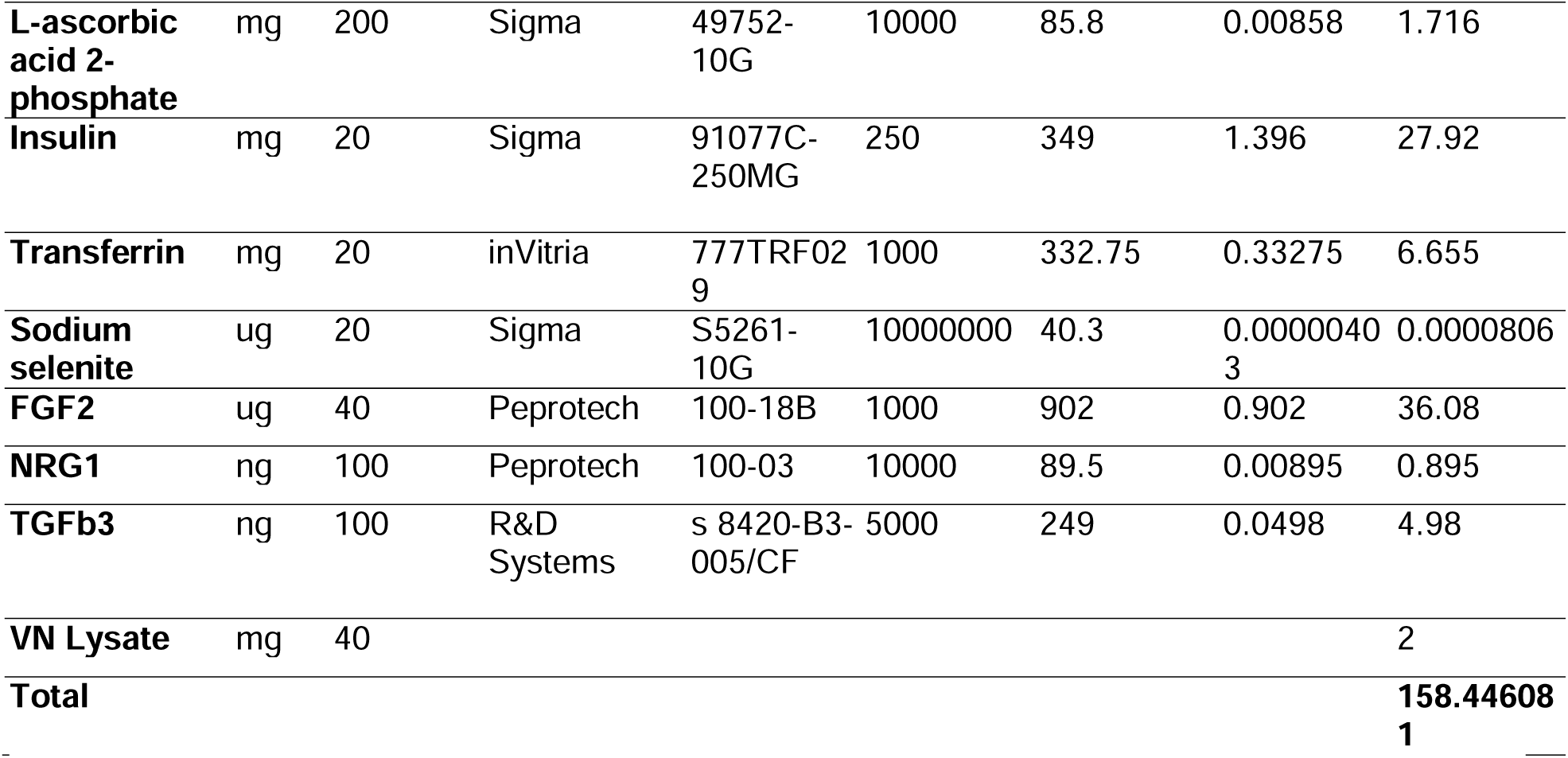
Cost comparison of itemized media components and total cost of iBSCGM and B9 with VN40 medium. The cost calculation for VN lysate was based on LB media (BD Difco, 244620). One gram of LB costs 0.346 USD. We prepare 25 mL of VN culture in LB, yielding approximately 4 mg/mL of lysate (total protein content). For 25 mL of culture, we use 0.625 grams of LB. Therefore, 0.625 grams of LB yields 4 mg/mL of lysate, and 40 µg/mL lysate is produced from 0.006 grams of LB. The cost of 0.006 grams of LB is $0.002, making the cost for 40 µg/mL VN lysate $0.002. This translates to $0.00005 per 1 µg/mL of VN lysate, and thus, 40,000 µg/mL would cost ∼$2.

**Table S3:**
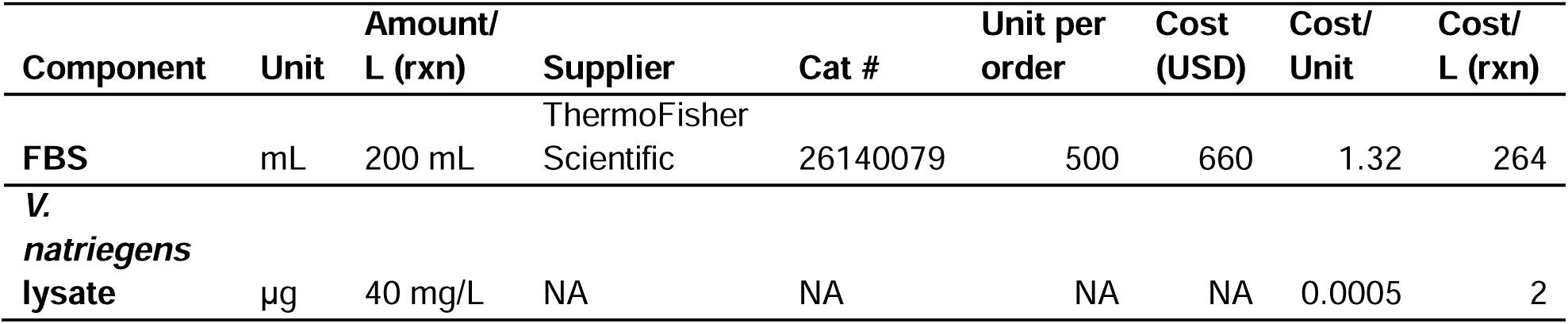
VN lysate is 99.2% cheaper than FBS.

### Additional details on LC-MS/MS methods

#### Reverse phase (RP) chromatography method

(Phenomenex Synergi Hydro-RP Column)

- Solvents: A: 0.1% formic acid in water B: 0.1% formic acid in methanol
- Column temperature: 15°C
- Flow rate: 0.2 mL/min
- Ion source: Turbo spray (ESI)
- Ion source Gas 1: 35
- Ion source Gas 2: 45
- Curtain Gas: 25
- Temperature: 450C
- IonSpray Voltage Floating: ±4500 V

Gradient:

| Time (min) | %B |
| --- | --- |
| 0-8 | 3 |
| 8-38 | 3 → 95 |
| 38-45 | 95 |
| 45-47 | 95 → 3 |
| 47-55 | 3 |

#### Hydrophilic interaction chromatography (HILIC) method

(Phenomenex Luna NH2 Column)

- Solvents A: 95:5 water:acetonitrile

+ 20mM ammonium acetate, pH to 9.45 using ammonium hydroxide B: 100% acetonitrile
- Column temperature: 25°C
- Ion source: Turbo spray (ESI)
- Ion source Gas 1: 35
- Ion source Gas 2: 45
- Curtain Gas: 25
- Temperature: 450C
- IonSpray Voltage Floating: ±5500 V

Gradient:

| Time (min) | %B |
| --- | --- |
| 0-15 | 85 → 0 |
| 15-28 | 0 |
| 28-30 | 0 → 85 |
| 30-60 | 85 |

